# Conceptual content in images triggers rapid shifts of covert attention

**DOI:** 10.1101/259929

**Authors:** Brad Wyble, Michael Hess, Chloe Callahan-Flintoft, Charles Folk

## Abstract

The visual system can use conceptual information to search for targets even in the absence of clear featural signifiers^1^, and visual saccades are often directed at target objects defined by conceptual content^2^. These abilities are a core component of our facility with the visual world. Here, we evaluate whether contingent mechanisms of visual attention, known to trigger in response to target features such as motion, color or luminance^3^, are also triggered by visual patterns that match conceptually specified categories. These pre-registered experiments provide convergent behavioral and electrophysiological support that covert spatial attention is rapidly triggered by natural image exemplars from superordinate conceptually described target sets such as *dinner food* or *four-legged animal*, even when each target was viewed only once. In the behavioral experiment when two targets were presented with onsets separated by only 167ms, subjects reported the second target more often when it was in the same spatial location as the first. In the EEG experiment, images elicited clear N2pc and P3 components only when they matched the conceptually specified target set. The latency of the N2pc peaked at roughly 250ms, which is comparable to that commonly found in other N2pc studies for simpler stimulus types. These results suggest that vision quickly decodes conceptual information from natural images and selectively deploys spatial attention to locations containing information that matches current search goals.

## Introduction

Effective, real-time interaction with the visual world requires the efficient allocation of limited processing and memory resources to those stimuli most relevant to current goals. Finding a lost child in a crowd, for example, involves allocating spatial attention to the subset of people that share perceptual features with the child (e.g., small, blond hair, etc). Indeed, a large body of research reveals that top-down attentional guidance is based on configuring the attention allocation system for “simple” visual features, e.g., color, orientation, or ^4-5^, or the relationships between those features e.g., “bigger” or “redder”^6^. However, it is often the case that stimuli relevant to behavioral goals are not defined by specific perceptual features, but rather by relatively abstract semantic boundaries. Searching for “sports equipment” in a department store, for example, cannot be accomplished by setting the system for simple features, such as color, because of the heterogeneity of featural content in the category exemplars and the fact that such features are generally not unique to that category. An important question, therefore, is whether the attentional system can rapidly decode conceptual information from natural images and use that information to guide attention. In this sense, *conceptual* refers to properties of a stimulus that relate to its placement in a semantic hierarchy^7^, that can be described verbally, and that do not rely on specific visual properties.

The groundbreaking work of Potter^1^ suggested that the visual system of the average adult is able to scan a rapid serial visual presentation (RSVP) of never-before-seen images in search of a particular image that matches a text descriptor such as *road with cars* or *girl sitting on bed*. Subjects were able to reliably detect a target image from a series of distractor images at rates of 113ms/image. Importantly, at this same rate of presentation, subjects were unable to non-selectively store memories of all the images in the sequence, resulting in very poor recognition accuracy for randomly probed images. Taken together, these results suggest that the visual system was able to select images that matched a conceptually specified descriptor. This result exemplifies one of the most mysterious and powerful aspects of perception, which is the ability for goals to change the selectivity of processing in favor of task relevant information specified at a conceptual level. Furthermore, while there was no specific measure of attention, the work suggested that the specification of attentional filters can happen at a very high level, using verbal descriptors that specified semantic properties.

An important step in understanding how attention is guided by goals was taken by Folk, Remington and Johnston^3^ who found evidence that task set influences which stimuli are able to capture spatial attention when subjects were looking for targets defined by features such as motion or color. In their experiments, the mere presence of a distractor containing the task-relevant feature prior to a search array made subjects slightly slower to report a target presented at a different spatial location, and faster to report at the same location. The same distractor had no such effect when its features were task-irrelevant. These findings suggest that goals can alter the ability of particular features to trigger attention. However, the extent to which this occurs for conceptual signifiers remains unknown. The work of Potter^1^ is suggestive of extremely rapid, conceptually driven triggering of attention, but does not speak to spatial attention as a mechanism.

Rapid triggering of attention by conceptual attributes of natural images would be difficult to explain using conventional theories of attention in which low-level features are used for filtering^4-5^ and attention binds features at each location in an image for more complex analysis. Moreover, single unit data from monkeys^8-9^ suggests that the visual system is adept at extracting high-level conceptual and categorical information from a visual stimulus. If the output of late-stages of the visual system can interact with top-down goal settings then it should be possible for attentional filtering to keep pace with RSVP streams.

Another source of evidence for the ability of conceptual information to directly trigger spatial attention is that natural images containing emotional or threatening content can capture attention independently of the featural content^10-11^. In addition, eye movement studies have shown that the eyes are drawn to exemplars that are semantically related but featurally dissimilar to target exemplars, e.g., fixating on a motorcycle helmet when searching for a motorcycle^12^. Verbal descriptors have also been found to elicit spontaneous capture of attention by line drawings that match^13^.

More recently, using an RSVP task,^14^ found evidence that peripherally presented distractors consisting of natural image exemplars of superordinate semantic categories can impair the detection/identification of centrally presented target exemplars if the distractors are members of the same superordinate category as the target. Notably, the very same distractors had no effect when the target was from a different superordinate category, ruling out salience-based distraction. It is unclear, however, whether these results reflect the misallocation of spatial attention, or more “central” processes, such as competition in visual short-term working memory. Another source of evidence that conceptually specified images can trigger attention is a finding of neural correlates of attention in response to line drawings that match a target set like items of clothing or kitchen objects^15^. While this result directly implies a role of attention, subjects see the targets many times, which might permit learning of specific exemplars. To test if spatial attention can be triggered by images that match a participant’s conceptually specified target set, we ran a pair of studies that attempted to link behavioral correlates of target detection performance to two neural signatures of visual target processing: the N2pc and the P3 components. The first experiment provided a behavioral measure of whether a target image selectively affected the processing of information at its location as predicted by a spatial attention account. The second experiment compared the N2pc and P3 components elicited by a target image with those elicited by control images that were exactly matched for content across participants. The N2pc is typically associated with the deployment of spatial attention^16-17^, while the posterior P3 (or P300 as it sometimes termed) is typically associated with the updating of working memory^18^. Together, these potentials would provide an indication that a target image is able to trigger attention and also be stored in memory. Moreover, by using a sufficiently large battery of images we can ensure that each target image is shown only once per subject which precludes familiarity with exemplars as an explanation for the observed results.

In both experiments subjects viewed two simultaneous RSVP streams of natural images, in search of objects belonging to one of 16 superordinate categories, such as *dinner food, four-legged animal, weapon, etc*. Subjects typed in the name of the target(s) they saw at the end of each trial (e.g., *hot dog, elephant, pistol)*. Target images were used only once per subject in both experiments.

The first experiment focused on behavioral correlates of attention. Subjects observed one or two targets per trial while varying whether the two targets were in the same or different streams. If targets are able to trigger a spatially selective attentional shift, then a second target in the same location as a first should be reported more accurately compared to a target in a different stream, which has been shown using simpler categorical stimuli^19-20^.

The second experiment recorded EEG while subjects looked for one target. Unbeknownst to the subject, each trial had a false target in it, which was an image that would be used as a target for another subject, but did not match this particular subject’s task-set at any point during the experiment. This manipulation ensured that ERPs elicited by the target were due solely to the match between the target image and the subject’s attentional set. If the images captured attention due to their intrinsic salience or content, then the effects would also occur when those images did not match the target set of the subject. Both target and false target images were only presented once to each subject.

Finally, a joint analysis of the two experiments, which used two distinct groups of subjects, measured whether behavioral accuracy in the first experiment for each image category was correlated with the neural correlates of attention and memory for the same set of categories in the second experiment. The use of a twin experimental design ensured that any correspondence between the behavioral and EEG results was not due to circumstantial variability in attention over time.

## Behavioral Data - Experiment 1 Results

Experiment 1 tested whether conceptually specified target images affect the spatial distribution of attention. In some of the trials, the two images, T1 and T2 were in the same RSVP stream, while in other trials they were presented in the RSVP stream on the opposite side of the display. Figure 1 illustrates a comparison between three conditions: Tl-alone, T2-same stream, and T2-different stream. This analysis plan, preregistered at https://osf.io/gdzxv/, made one key prediction and four secondary predictions. The key prediction was that T2 accuracy will be significantly higher in the same stream condition than in the different stream condition 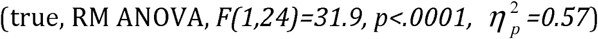. This prediction was key because it was the clearest comparison of similar target conditions, varying only the spatial similarity of T1 and T2. Our four secondary predictions were that T2 accuracy would be higher in the same-stream condition compared to T1 in the same-stream condition because the T1 recruits attention which benefits the T2 condition 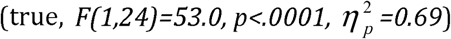; T1 accuracy would be higher in the different-stream condition compared to T2 in the different-stream condition because the attention is specific to the T1 location 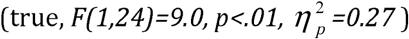; T2 accuracy would be higher in the same-stream condition than T1 alone because of the same attentional effect 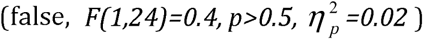 and finally that T2 accuracy in the different stream condition would be lower than T1 alone, because T2 suffers from attention between triggered by the Tl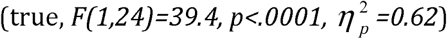.

**Figure 1.**
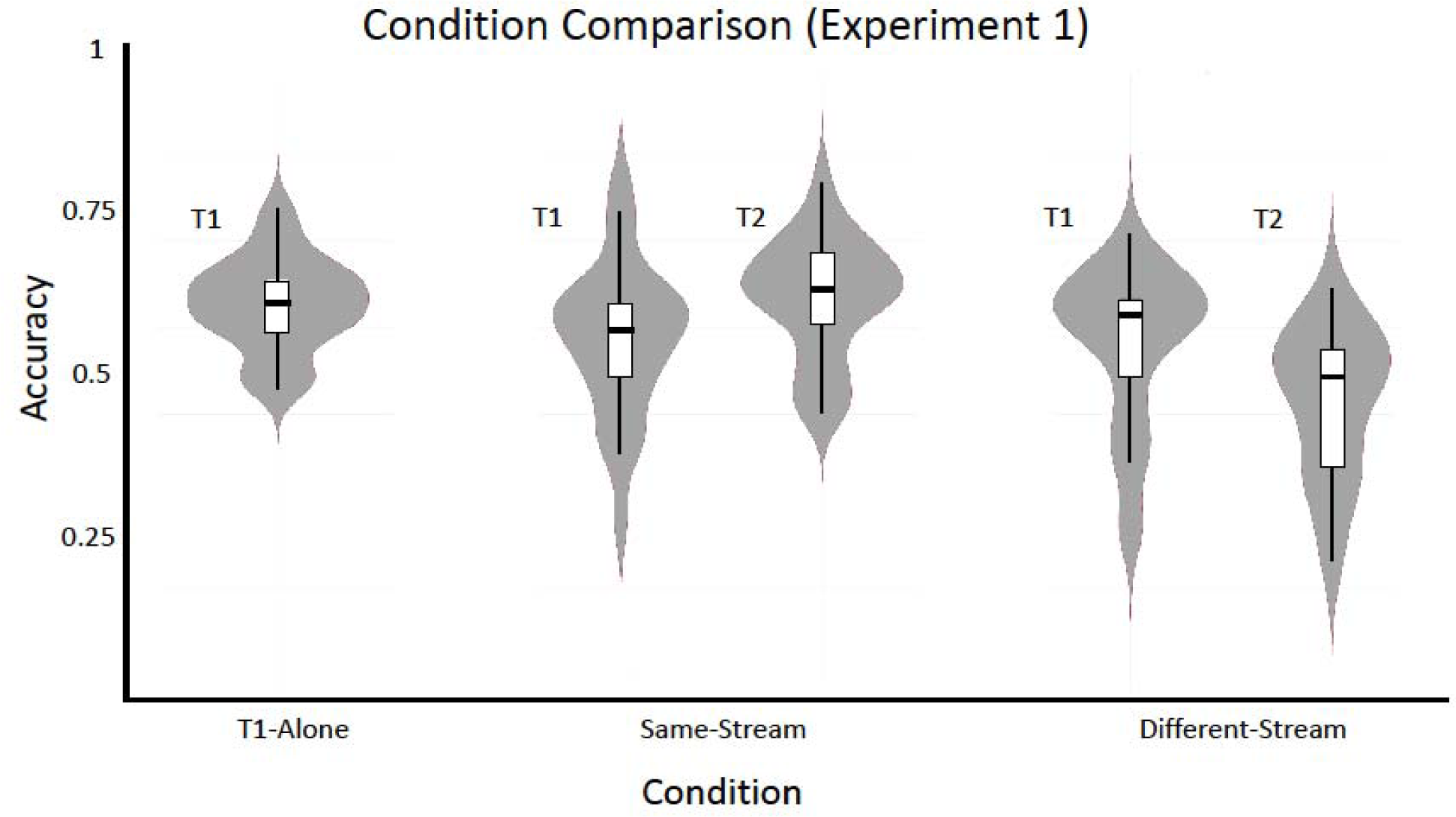
A violin plot of the behavioral data distributed by condition. When two targets were presented in the same stream, T2 was reported more often than when the two targets were presented in the opposite streams.

These behavioral results support an account in which an image matching a target category would trigger spatially selective attention. The key prediction, which is that T2 would be better in the same compared to the different stream as the T1 revealed a strong effect. Of the other 4 secondary predictions, 3 were significant in the predicted direction. The failed comparison was the most ambitious, as it predicted T2 accuracy in the T1 stream would be higher than the accuracy of detecting a solitary target. The failure of this prediction suggests that the interference caused by having two targets in succession provided a counterweight to the benefit of spatial selection for the T2. As a result, T2 accuracy in the same-stream condition was approximately equal to report of a single target rather than being higher

Experiment 2 was intended to test for convergent ERP support that the target image was able to trigger spatial attention as well as test for evidence that he target was encoded into memory subsequent to attention

## EEG Data - Experiment 2 Results (one target per trial)

This experiment was similar to Experiment 1 except that there was one target and one false target in each trial, presented at different times. On average, subjects correctly reported the target 59% (+/- 2.14%) of the time.

For the ERPs, a preregistered analysis plan, using a cluster-mass approach^21^, linked here: https://osf.io/84rvy/. predicted that there would be an N2pc for the true targets but not the false targets, with a peak latency at nearly 250ms, and a maximum in posterior parietal electrodes. This prediction was confirmed by a confirmatory analysis, showing that targets elicited two significant cluster masses when contralateral and ipsilateral waveforms were compared (Figure 2a). The first significant cluster had a range of 236ms to 268 ms and was centered on electrodes 01, 02, P7 and P8 with FWER corrected p-values ranging from .022 to .046. The second significant cluster ranged from 276ms to 304ms on the P7, P8 electrodes, with FWER corrected p-values ranging from .03 to .049. A second confirmatory analysis revealed that false targets did not produce a similar significant cluster.

**Figure 2.**
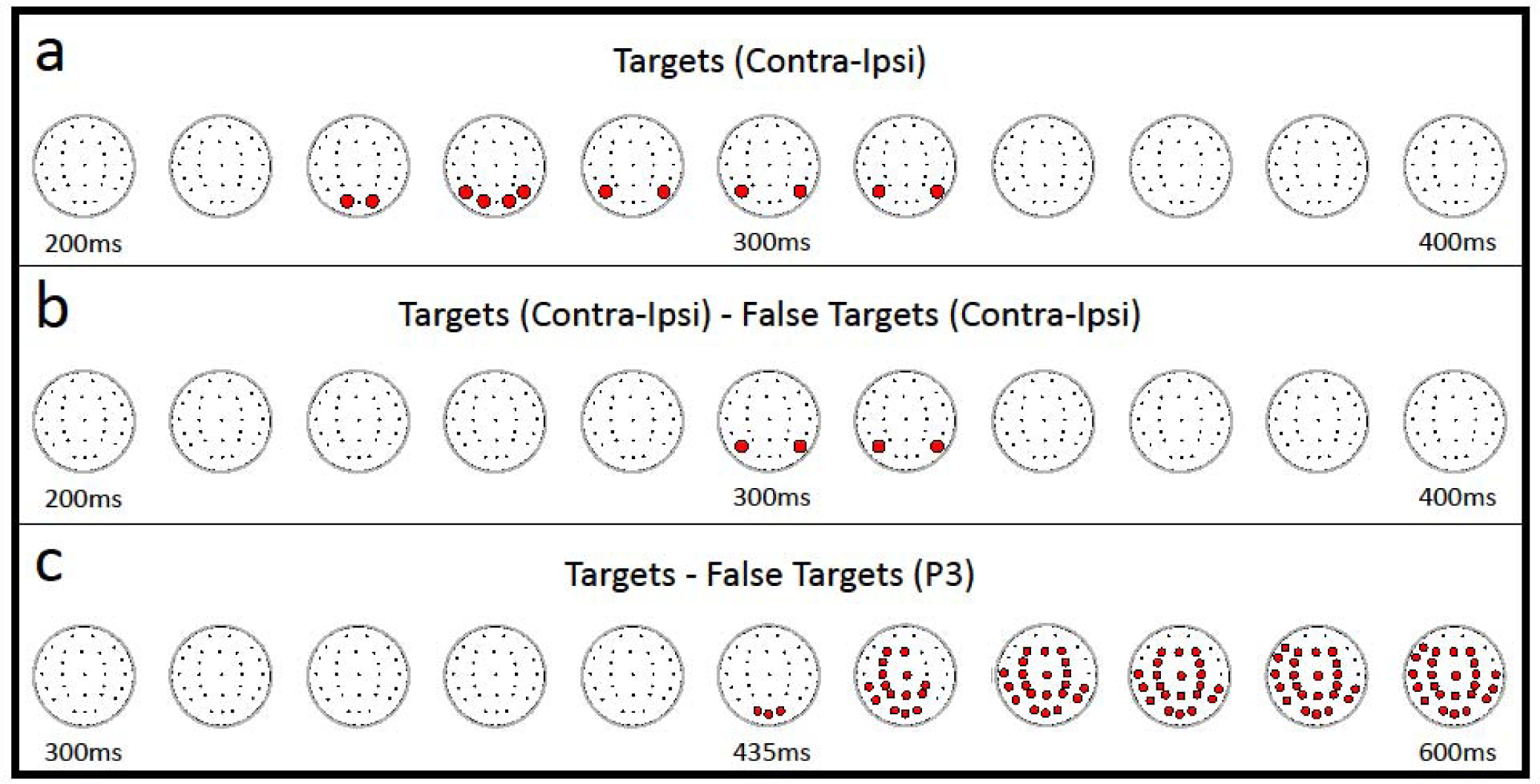
Visualizations of the cluster mass analyses. The cluster mass analysis uses a permutation to identify significant clusters of t-values across electrodes and time. Time (shown in milliseconds) is relative to the onset of the stimulus. Each dot represents an electrode, with red dots indicating membership in the contiguous cluster in this window at each time 20ms bin. All scalp maps are oriented on the axial plane so that the top represents the front of the head, **(a)** Two significant clusters from comparing contralateral to ipsilateral waveforms in relation to the targets in the 200-400ms window (the short gap between the clusters is not visible at this temporal resolution), **(b)** Significant cluster when comparing the contra-ipsi difference waves of targets and false targets in the 200-400ms window, **(c)** Significant cluster when comparing targets and false targets in the 300ms to 600ms window across all electrodes without regard to target laterality.

**Figure 3.**
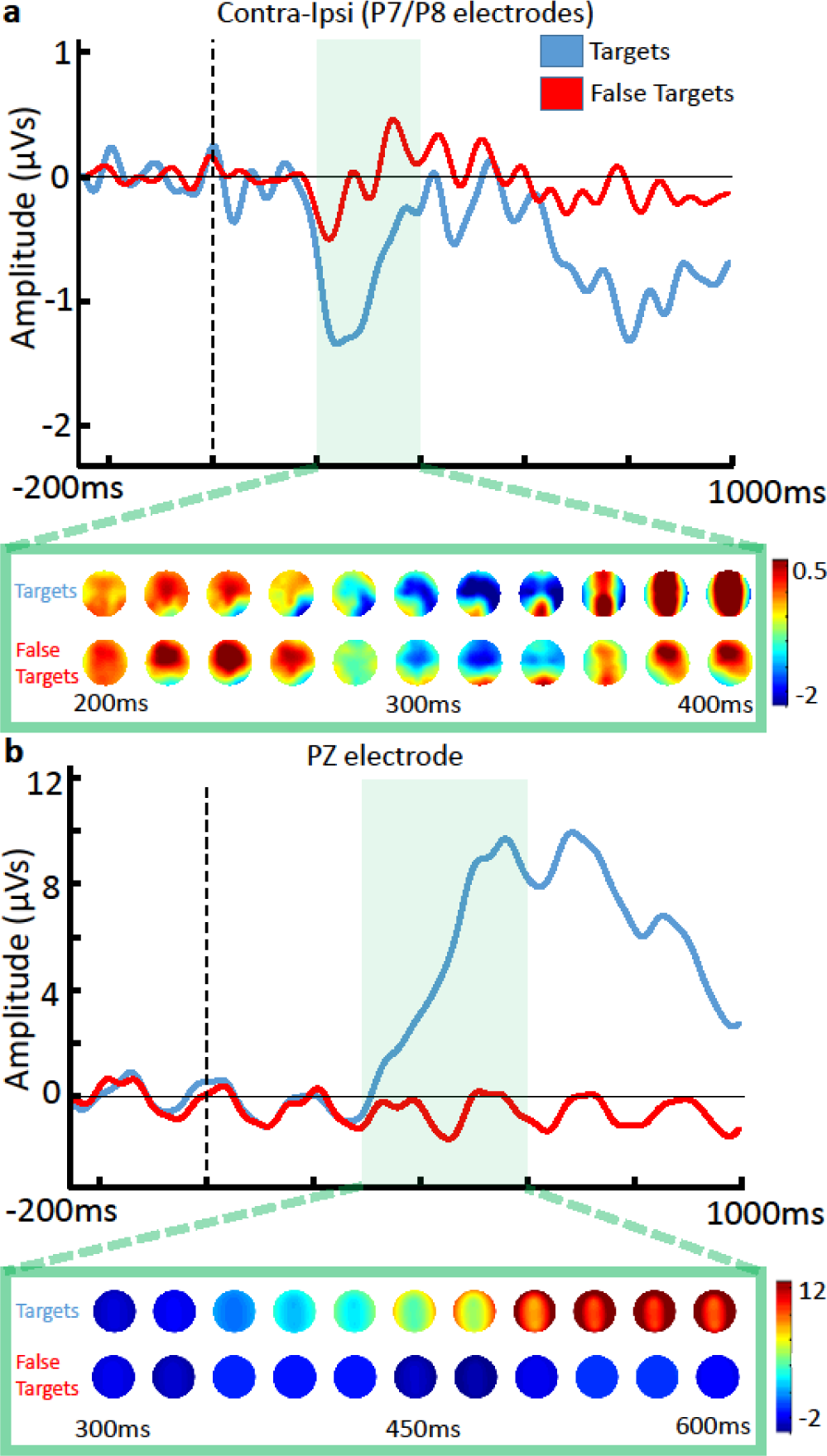
ERPs and scalp maps illustrating N2pc **(a)** and P3 **(b)** components for targets and false targets. Scalp maps indicate the distribution of voltages for the predefined time window used in the cluster mass analysis. Scalp maps are corrected for target location by left/right reversal of electrode locations for trials in which the target was on the right. **(a)** The N2pc was calculated by subtracting the contralateral from the ipsilateral of the P7 and P8 electrode pair. A visible indication of the N2pc in the contralateral (right) side of the scalp begins at the 240ms map for targets and remains up to the 320ms plot, **(b)** The ERP output of the PZ electrode is selected to show the P3 component.

Furthermore, an exploratory comparison that was not pre-registered compared the contra-ipsi difference waves from target and false target trials. It revealed a significant cluster mass peaking at 300ms on electrodes P7 and P8 (FWER corrected *p* < .05 in 4ms timesteps from 288ms to 316ms, *p* = .005 at peak) (Figure 2b).

A further exploratory analysis looked for evidence of a target-evoked P3 component, which would suggest that working memory was updated by the contents of the target image. As seen in Figure 2c, a comparison of target to false targets revealed a highly significant cluster mass, roughly in the time range 300-600ms, peaking at electrode PZ, but broadly distributed across the scalp (FWER corrected *p* < .05 for 4ms timesteps from 432ms to the end of the time window, *p* < .0001 at peak).

These results indicate that perceiving a conceptually specified target evokes neural correlates (the N2pc and P3) that resemble the components typically associated with attention and memory updating for targets specified by simple designators such as color or luminance, more complex designators such as the categorical identity of a word or letter, and ink drawings^15-17,20^. In this case, these correlates are not triggered by an intrinsic property of the images, since those same images evoked neither an N2pc nor a P3 when they were used as false targets for other subjects. To mirror more conventional ERP analyses, a window-based analysis was also used to compare the average voltage in the N2pc time window (200ms-400ms) for the contralateral difference wave of electrodes, P7 and P8 between targets and false targets, revealing a statistically significant difference 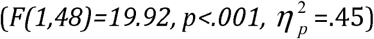. Similarly, a significant difference was found between the target and false target waveforms from electrode Pz within the P3 window i.e. 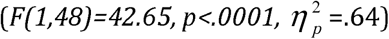.

To determine if the false targets played a role in the failure to elicit these correlates, a cluster mass analysis was performed on the comparison between contralateral and ipsilateral electrodes for false targets in the 200ms-400ms window, only including trials in which the participants responded *incorrectly* to the identity of the target. This analysis revealed no significant indication of an N2pc either (p=0.29). Additionally, a cluster mass comparison analysis was performed using these “incorrect” trials as well as the false target data for all trials, and no significant difference was found between the two (*p* > = 0.79). Thus, there is no strong evidence that false targets triggered attention or were encoded into memory.

## Median split analysis

To determine whether images can trigger attention by virtue of their content and that this form of attention facilitates the ability to perceive and remember target images, as in Potter^1^, a follow-up analysis compared behavioral accuracy of images to the N2pcs across experiments. The logic of this comparison is that if attention plays a role in perceiving these targets, then behavioral accuracy should be correlated with their ability to trigger attention on a superordinate level, as indexed by the N2pc. Importantly, this comparison was done across-experiment, which ensures that any observed effect is not due to fluctuations in attention over time. The analysis began with a median split of the 16 categories according to the average accuracy of reporting those targets in the T1-alone condition of Experiment 1. We then applied this same division to the categories from Experiment 2 and compared both accuracy and the N2pc from the best-8 categories to the worst-8 categories (Figure 4). In terms of accuracy there was a strong correspondence between the two experiments, with 7 of the 8 categories being in the same half of the median split.

**Figure 4.**
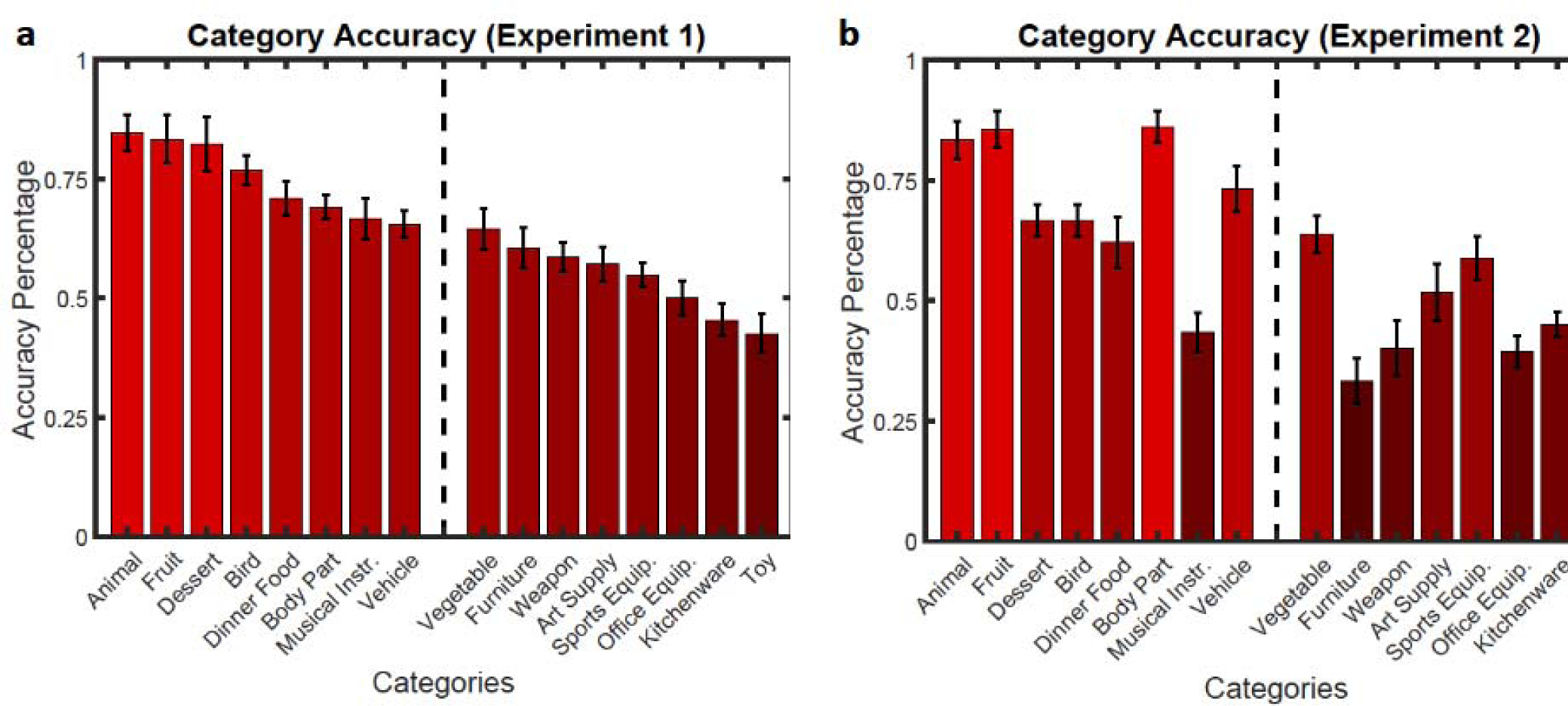
The mean category accuracies for the behavioral **(a)** and EEG **(b)** datasets, ordered from Experiment l’s most to least accurate. This ordering allows a visual representation of the similarity in the categories’ relative accuracy values between experiments. The dashed line indicates the median split taken from the behavioral dataset. This split was used to group data in the EEG dataset to determine if there is a significant difference in the N2pc time window (200ms-400ms) between the two groups.

A comparison of the difference waveforms associated with the top 8 and bottom 8 trials along with topographic plots are shown in Figure 5. To determine if there was a link between the neural correlates of attention and the accuracy of reporting the different kinds of stimuli, an exploratory cluster mass comparison analysis of the contra-ipsi difference waves for correct report trials revealed that a cluster corresponding to the N2pc was larger in the high-accuracy group compared to the low-accuracy group (*p* < .01). A cluster mass comparison analysis of the P3 window however revealed no significant clusters (*p* > = 0.16). This suggests that the target categories in the lower half of the median split for accuracy in experiment 1 were also less effective at triggering attention in experiment 2, as indexed by the N2pc. As it was possible for this result to be an artifact of differences in trial count between the top and bottom halves of the category sets, a bootstrap analysis in the supplemental section equalized trial count in the two halves and still found a difference in N2pc amplitude between the conditions.

**Figure 5.**
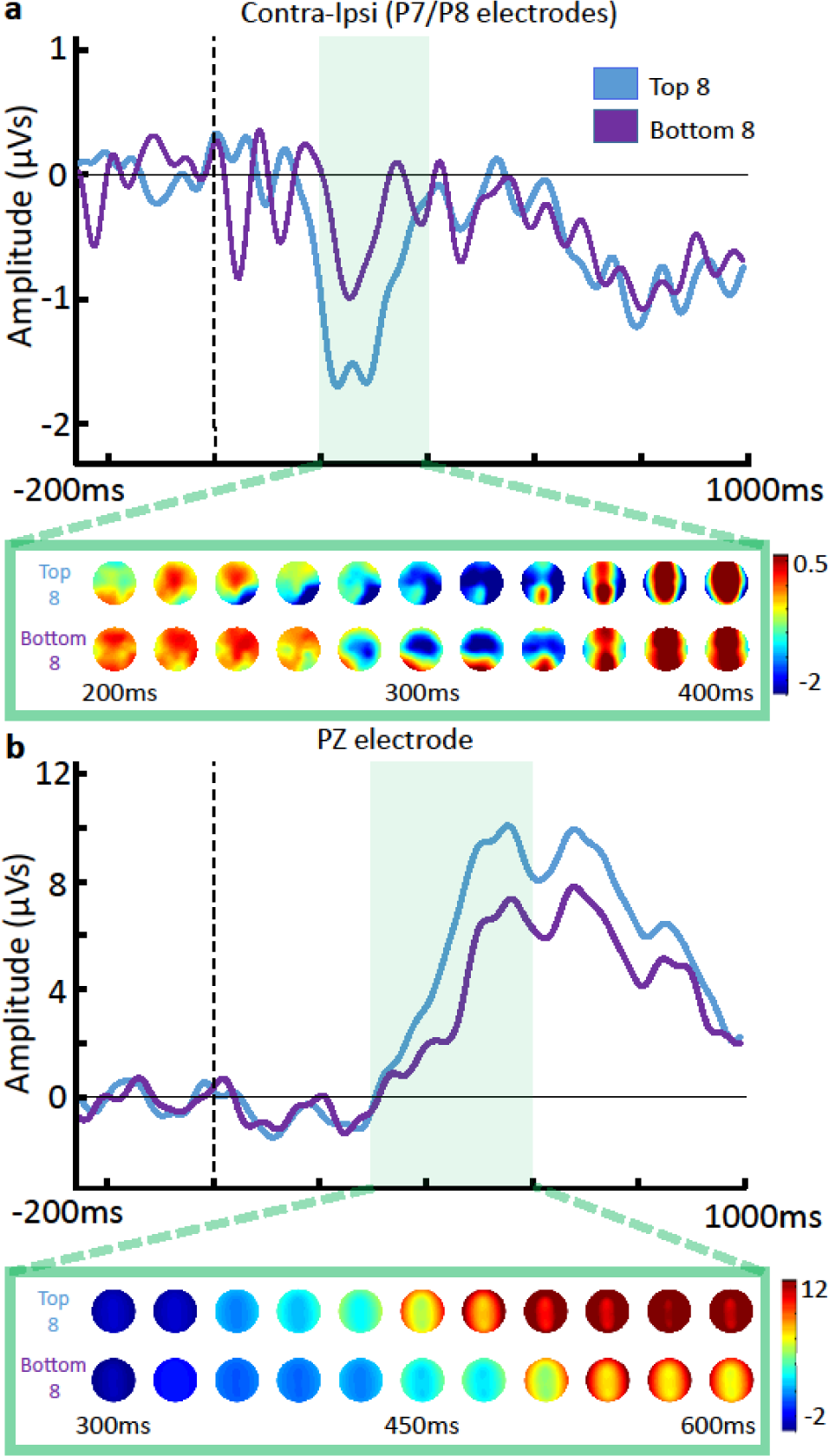
ERPs illustrating N2pc **(a)** and P3 **(b)** components for the categories median-split by accuracy. Below each ERP plot are the ERP amplitudes projected onto topoplots, with the left side of the head representing ipsilaterality in relation to the target or false target, and the right side representing contralaterality.

## Discussion

These experiments provide convergent behavioral and neural evidence that the visual system is able to decode conceptual information from a rapid presentation of natural images that had not been previously viewed. That computation’s result could trigger a spatially selective form of attention when an image matched a participant’s search goal, specified at a superordinate categorical level. Furthermore, a false-target control demonstrated that the triggering of the ERP correlate of attention was not caused by the inherent physical properties of the target images, but instead was a property of the image’s match with the participant’s current goals. These results extend the scope of the contingent orienting hypothesis to include conceptual information, in addition to the simpler features such as motion and color that have traditionally been studied. It is important to note, however, that although the present results provide clear evidence for contingent shifts of spatial attention to conceptually relevant stimuli, it is unclear whether these shifts of attention represent attentional *capture*. Evidence of capture would require similar effects when the eliciting stimulus is completely task irrelevant, which is not the case in the current studies(but see [14])

This finding suggests that rapid, feed-forward processing in the ventral stream^8,24^ is not just useful for identification, but can also be used to trigger spatially selective attention contingently when a natural image matches a conceptually specified superordinate attentional set on first exposure to that image. The data suggest that when someone is asked to find an unfamiliar object specified only by its category and not its features (i.e. find the hat on the table without foreknowledge of what the hat looks like), the visual system is able to rapidly deploy attention to the location of the visual field containing an exemplar of that category, even when it is outside of the fovea. This triggering of attention could play a role in enhancing the representation of the target object, and also driving visual saccades to the object’s location^2,24^.

The most surprising aspect of these findings is that the latency of the N2pc, considered an EEG marker of attentional engagement to visual targets defined by highly characteristic low-level visual features, such as colors, or repeated use of the same word^16-17^ is not greatly increased when targets are natural images that have never been seen before and participants are using a conceptually defined target set. The rapidity of attentional onset in this case argues against theories that focal attention is a necessary precursor to using conjunctions of features to identify higher-order regularities to decode conceptual information in a natural image. These findings are in agreement with VanRullen^25^ who suggests that visual experience ingrains the ventral pathway with “hardwired” bindings of features for familiar visual stimuli and these hardwired bindings permit them to be discriminated without focal attention. Such a system would be able to provide a rapid, initial assessment of the conceptual content of each item, and deploy attention selectively to specific items that match a conceptually designed target set. In contrast, tasks that require conjunctions of features that do not map onto well learned, meaningful representations, would be impossible at that speed. Thus, finding a specific hànzì character in an RSVP at 150ms is trivial for someone literate in Chinese but extremely difficult for one who is not.

Computer vision algorithms equipped with deep learning architectures^26-27^ are increasingly playing a role in understanding how such hardwired computations could be performed by the human visual system. Such architectures reveal that a succession of processing layers in a neural network is able to map natural images onto lower-dimensional representations that can be classified by labels that a typical human observer would apply. During the image decoding process, such networks operate in a purely feedforward fashion and have operators that are neurally plausible at each step. As discussed below, it would be straightforward to use such representations to trigger attention selectively at a spatial location.

## Attentional set for categories

The rapid decoding of the content of each image can be coupled with a form of attentional set, governed by goals or task instructions. It has been shown that asking subjects to look for particular kinds of objects, such as vehicles or people in a natural scene, induces changes in cortical activation that are suggestive of tuning of visual processing towards those categories. Some of these changes are clustered in areas late in the ventral stream^28-29^ and others have been found throughout much of the cortex^30^. These changes are consistent with the idea that the visual system has a ‘what’ template, that can be configured so that certain brain areas respond selectively to information that matches a specific template^31^.

From such a template, it would be straightforward to extract a coarse-grained spatial signal to indicate what part of the visual field contains the matching information. For example, neurons near the anterior end of the ventral visual pathway, which combine high-specificity pattern detection with coarse-grained spatiotopy^32^, could provide input to higher order networks trained to detect conceptually meaningful patterns. In computer vision, such classification is accomplished with tools such as support vector machines and neural network classifiers that decode representations from the upper tiers of a feedforward network^26,33^. If a neural equivalent of such classifiers were coarsely tiled across the visual field (Figure 6a), their output could indicate not just the presence, but also the general location of information matching the attentional set. The visual system would detect a match between the top-down specification of a concept, and stimulus-driven activation of these concept detection networks. In response to such a match, attention could be triggered selectively at the location of the visual field corresponding to the activated portion of the network. The time course of such feed-forward computation could be extremely rapid according to single unit data from monkeys. For example, the identity and category of a complex visual stimulus can be decoded within 125ms of stimulus onset in macaque IT neurons^9^. The active deployment of attention could be only several synapses beyond such evoked activity. Such a rapid attentional response would explain the spatial selectivity of attending to the second image in Experiment 1 here, as well as the spatially selective changes in reaction times in response to a dot probe appearing just 117ms after a car or person silhouette when viewers are searching for cars or people, respectively^34^. These results dovetail with complementary results from MEG, showing that information related to target category can be retrieved from posterior brain regions to a superior degree relative to distractors starting in a brief window 180-220ms after image onset^35^.

**Figure 6.**
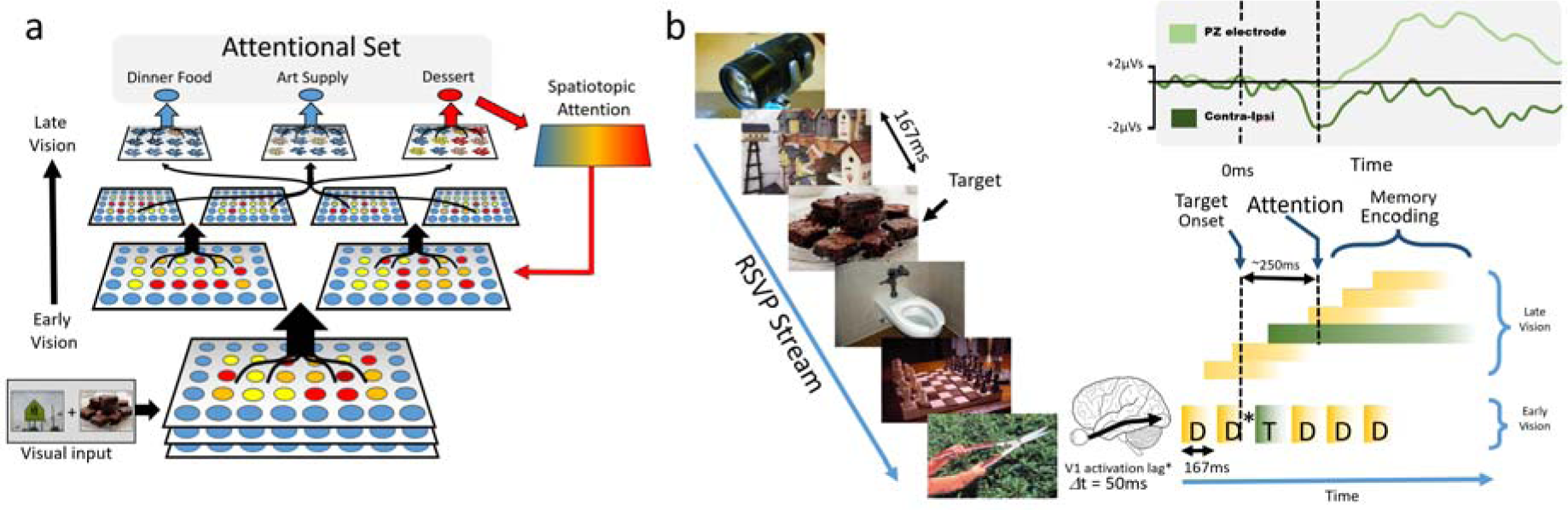
**(a)** Illustration of how a hierarchical feature detector, would be able to rapidly map visual images onto conceptual representations that could be used to drive classifiers. In computer vision, models with architectures similar to these have been able to correctly classify natural images into categories. By tiling such classification across a population of cells with coarse spatial specificity, the model could rapidly determine the category of an image and also its general location within the visual field. Stimuli that match a top-down attentional set would be able to trigger a spatially selective deployment of attention, **(b)** Illustration of how contingent attention with a latency of 250ms would be able to select information from RSVP at 167ms RSVP. Attention, triggered by a match between an image and a conceptual target set would become active at ~200ms after a stimulus representation reaches visual cortex (there is a roughly 50ms latency between retinal and visual cortex activation^39^). Given that higher-level representations in the ventral stream will be lagged relative to their lower-level sources during RSVP^9^ attention at this latency would still have time to act on the delayed representations at higher levels of processing. The ERP signals reported here (top right corner) are from experiment 2 and provide a metric of the time course of attention (N2pc, dark green trace) and encoding processes (P3, light green trace)within the visual pipeline, which is illustrated in hypothetical form below.

### On the latency of attention

Returning to the starting point of this investigation, the initial hypothesis was that the deployment of attention aids in the detection of target images in RSVP paradigms. One apparent difficulty with this explanation is that our measure of the latency of attention, using the N2pc, puts the onset at approximately 250ms post stimulus. However, in RSVP, stimuli often arrive so quickly that the representation of the target in early visual areas would be replaced by that of the following image well before 250ms had elapsed. Thus, it seems that attention would onset too late to enhance the representation of the target image which triggered it in the first place. This discrepancy can be resolved by considering that the visual system operates with a pipelined architecture, in which deeper layers of processing are time lagged relative to earlier layers, and also exhibit less strict temporal masking effects^8^. Figure 6b illustrates a conceptualization of how a series of images presented at 167ms could evoke a succession of representations in higher level visual areas. In this framework, attention might act on those higher level representations, rather than the sensory trace in earlier visual areas. Thus, the encoded information might miss some of the fine-grained detail in a target image, while enhancing the conceptual information that is necessary to categorize and locate the stimulus. Recent work^36^ provides neural evidence of the pipelined nature of visual processing during RSVP, and also the prolonged processing of target information relative to distractors, as depicted in Figure 6b.

### On transient attention

Another question that arises when considering how visual attention might operate on a central RSVP paradigm is that spatial selection seems irrelevant when there is only one location. However, the phenomenon of ***transient*** attention^37^ suggests that spatial attention has a temporally selective component as well. Thus, even when subjects are focusing their voluntary spatial attentional resources toward a central RSVP stream, a transient attentional enhancement would be additive to the voluntary, sustained form of attention. Moreover, this transient form of attention is selective in space as well as time^38^, which makes it effective in rapid attentional selection regardless of whether one or multiple locations are being monitored.

In conclusion, the present findings indicate the speed and flexibility of contingent visual orienting by suggesting that even never-before-seen natural images will trigger the deployment of attention if they match an attentional set defined at the superordinate level. This capability is presumably a key factor in our ability to search for complex objects in real-world tasks, even in the absence of specific knowledge about the specific features of a particular object.

## Experiment1: Behavior

### Methods

#### Participants

Twenty-five students from the Pennsylvania State University research participation pool participated in this experiment in exchange for credit as a course requirement. This sample size was determined in advance of data collection according to experience with pilot studies.

#### Apparatus

The experiment was performed using Matlab 2012b with PsychToolbox-3^40^ extensions on a PC using a CRT monitor with a refresh rate of 75Hz at 1024 x 768 resolution.

#### Stimuli

The stimuli were partly taken from previous studies^14,41^, with others downloaded from Google searches. The images consisted of colored photographs of single objects in their natural settings and pictured scenes containing multiple objects in their respective contextual settings. There were 384 target images and 653 distractor images in total, with an additional set of 40 images used in the practice trials. The distractor images consisted of single objects, multiple objects, scenery, animals, or people. The target images were organized into 16 categories of 24 images each. These categories included: Art supply, Bird, Body part, Dessert, Dinner food, Four-footed animal, Fruit, Furniture, Kitchenware, Musical instrument, Office equipment, Sports equipment, Toy, Vegetable, Vehicle, and Weapon. Each target set contained two exemplars of 12 different kinds of object, e.g., two unique pictures of an elephant in the Four-footed animal set. All images were resized to a dimension of 300 x 200 pixels. Images and their corresponding category folders can be found on an OSF page titled Category Behavioral RSVP, compressed in a folder named Stimuli here: https://osf.io/qbnxt/.

The 16 categories were randomly split into two groups of eight categories for each subject, one half were used for targets and the other half were used as additional distractors. Thus, each subject’s set of distractor images included distractors from the Distractor set as well as vetted images from the unused target sets. Each subject’s set of targets included 12 to 24 target images from each of the eight selected categories. The number of target images used in each block was contingent upon the number of trials in that block in which two targets were shown.

The eight target categories for each subject were presented within eight consecutive blocks of trials, one category per block. For each block, the distractor images were carefully vetted and excluded if they could have been easily mistaken for that block’s target set. On each trial, either one or two target images and either 47 or 46 distractors were shown in an RSVP stream. Each target image was seen only once per subject while distractors could be reused between trials but not within trials.

#### Procedure

Subjects searched through two parallel RSVP streams, each 24 items long for one or two targets. Except for two practice trials at the beginning of the experiment, the trials were organized into eight blocks, one for each target set, with the blocks randomly ordered for each subject. Each block contained 12 trials, with a total of 96 trials for all eight blocks. On each trial, the participant was given a superordinate category label (e.g., “Body Part”) and was asked to look for target images matching the label. Subjects pressed the spacebar to begin the trial at which point the fixation cross appeared, with the image RSVP streams beginning 267ms afterwards on the left and right of the cross. SOA between images in both streams was 167ms (except for the two practice trials and first two trials of the main experiment, see below) and there was no inter-stimulus interval (Figure 6). The entire length of the RSVP stream was 25 displays in 3.9 seconds. Targets could appear in a range of the 6th display to the 21st display. Two practice trials preceded the experiment. Practice trials were identical to the other trials except that they used unique target sets not present during the main experiment, i.e., Clothing and Plumbing, and images during these trials were presented at 817ms and 317ms SOA. The first two trials of the main experiment block were presented at 267ms and 217ms SOA respectively to help participants adapt to the rapid presentation rate. These two trials were excluded from analysis, as well as the practice trials.

There were three separate conditions which counterbalanced evenly across trials and were randomly ordered: *Single-Target*: only one target, *Same-Stream*: Two targets were shown in consecutive displays in the same RSVP stream, and *Different-Stream*: Two targets were shown in consecutive displays in opposite RSVP streams.

After the RSVP, the participants were asked “What is the first target object you saw?” to which they responded by typing in their answer without time pressure and then hitting Enter. Next they were asked, “What is the second target object you saw?” which they responded to the same way. Backspacing was allowed to correct responses and participants were made aware that they could type “nothing” to either response if they did not recall seeing the target(s).

**Figure 7.**
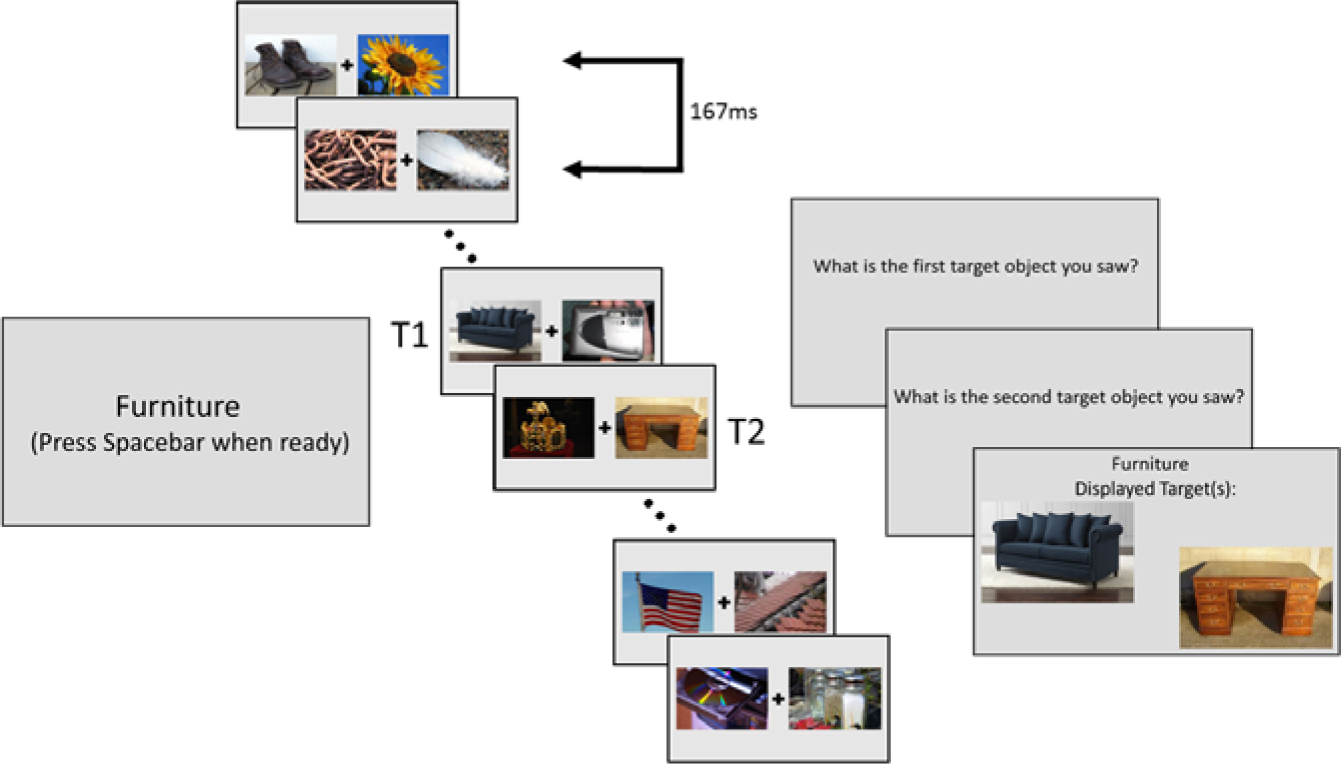
Layout of the *Different-Stream* condition in Experiment 1 (Note that the images are not to scale.) The category is given, then once the participant is ready, the spacebar is pressed and fixation cross appears on-screen. 217ms later, the RSVP streams begin. T1 and T2 on the figure indicate the presence of targets i.e. images that match the given category. Once the streams end, participants report the first and second target. They are then given feedback lasting one second which indicates which target images were shown. During feedback, the targets are displayed in the left or right position in which they were shown, with T1 presented slightly higher on screen than T2 to indicate ordering in the two target conditions (*Same-Stream* and *Different-Stream*.)

#### Scoring

Participants’ responses were scored by raters. Raters were presented with each target image shown during a trial as well as the corresponding category name and the participant’s response to the target. Raters responded “y” if they believed the participant responded correctly and “n” if they believed the participant responded incorrectly.

Volunteers were told to be fairly lenient when rating responses. For example, if a participant typed “round, yellow fruit” instead of “lemon,” the response would be marked as correct. Also, any obvious spelling errors were forgiven. Volunteers followed the guideline of “Do you believe, based on the participant’s response, that they saw this specific image?” in order to provide their ratings. Participant responses that were too general or that blatantly did not fit the target image (e.g., typing “rhino” when a tiger was shown) were marked as incorrect. The order of the subject’s two responses was randomly shuffled during scoring to prevent any possibility of bias on the part of the rater to reinforce the hypothesis.

## Experiment 2: EEG component

### Participants

Twenty-five students from the Pennsylvania State University research participation pool participated in this experiment in exchange for credit for a course requirement. This sample size was determined in advance according to a power analysis on data from a prior version of this experiment.

### Stimuli

The stimuli set was the same as in Experiment 1. The only addition was a symbol, either a period or a comma, which appeared at the end of the stream on 1/3 of the trials. In this paradigm, all targets from a randomly selected half of the categories were used providing eight blocks of 24 trials each. This compiles to a total of 192 trials, each with a unique target. The eight categories that were not selected as targets for a given subject were used to provide false targets, one per trial. False target categories are vetted on a category-by-category basis and only included if they contain no images that could be mistaken for a target in the current category. These false targets are presented on every trial and provide assurance that any target-specific ERP is due solely to the match between the target and the target-set given to the subject, rather than the categorical novelty or image statistics of the target, since those properties are similar between the target and false target image sets.

### Procedure

The instructions were the same as before, though participants were told that only one target would appear per trial, and that a period or comma may appear at the end of the stream which they would have to report as well. The period/comma task was used to encourage subjects to keep their eyes on the fixation cross until the end of the RSVP. The prompts were as follows: “What object did you see?” followed by “What symbol did you see?” with the latter only being asked if a symbol was shown. Again, the responses to these questions were typed in by the participant (Figure 8). Given the number of trials, breaks were inserted at the end of every block of trials to alleviate fatigue. The break would read, “You’ve earned a break! Press Spacebar when you’re ready to continue.”

**Figure 8.**
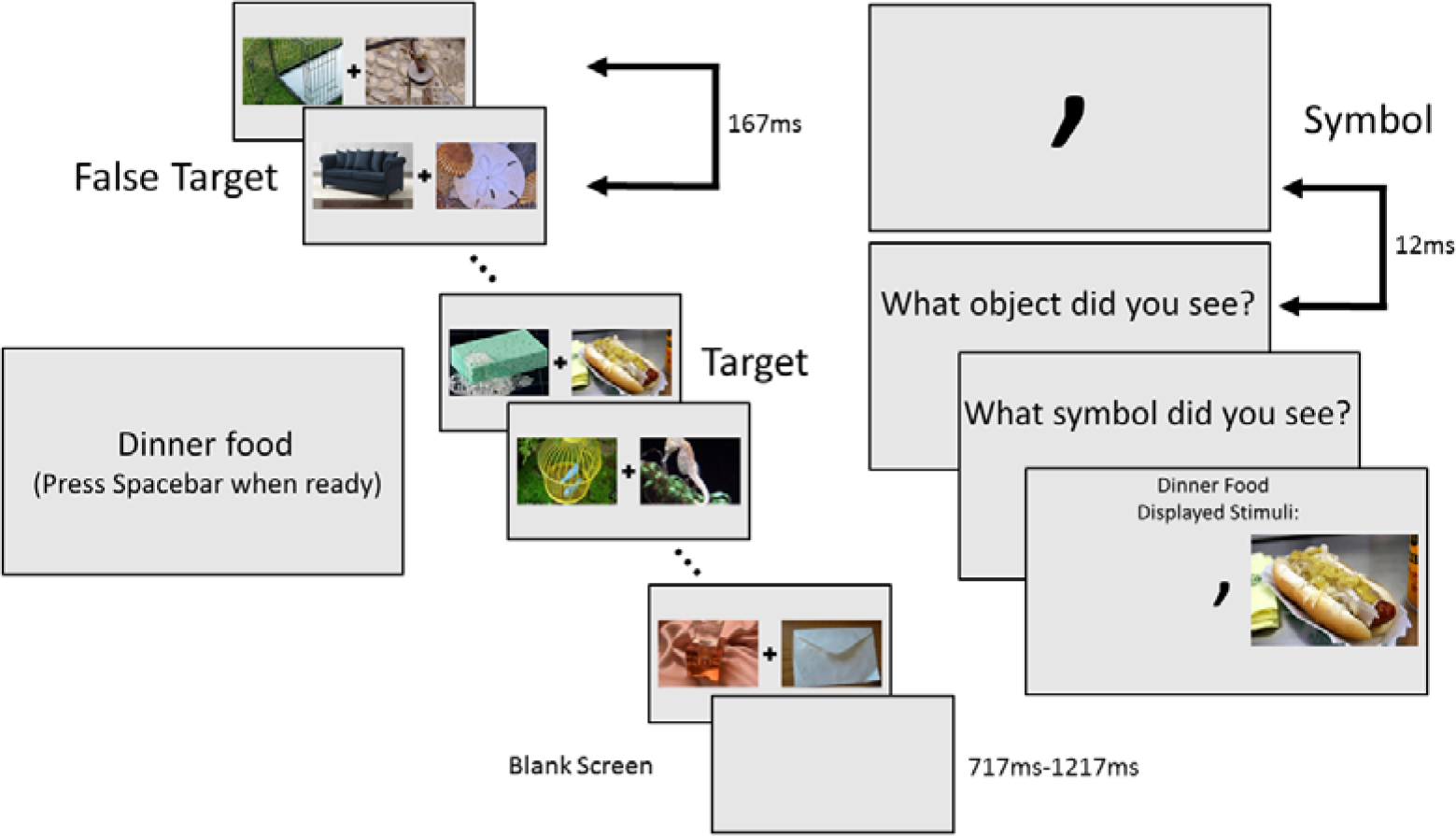
Layout of an Experiment 2 trial. Note that neither the images nor the symbol are to scale. The symbol was the same relative size as the fixation cross. If a symbol was to be shown, a jittered delay (717ms-1217ms) post-stream would occur, followed by the symbol which would appear at the center of the screen for 12ms.

SOA timing was exactly the same as Experiment 1 (167ms). The only difference in timing between the experiments is a jittered post RSVP stream delay (717ms-1217ms) that was followed by the presentation of a period or comma on 1/3 of the trials for 12ms. The stream length is the same as in the behavioral (25 displays), but the range of possible target display times is broken up into two possible ranges, either the 6th display to the 11th display or the 16th to the 21st display. The likelihood of the target displaying in either time range was 50%. The false target was placed into the other time range. This provided enough time between the target and false target display so that possible ERP components would not overlap. The stream containing the target (left or right) was randomly selected on every trial. The false target was then placed within the stream on the opposite side. This was done so that there was no possibility of conflating attentional effects towards the target and false target.

During analysis, the ERPs are time-locked to both the targets and false targets to detect if the selected target images have any special saliency or if attention deployment is truly driven by whether or not the image is a part of the target category.

### Apparatus

The experiment was run using Matlab 2012b with PsychToolbox-3 extensions on a PC using a CRT monitor with a refresh rate of 75Hz at 1024 x 768 resolution. EEG data were recorded by a Neuroscan EEG amplifier using a 32-channel sintered Ag/AgC1 electrode array mounted to an elastic cap according to the 10-20 system [FP1, FP2, F3, F4, FZ, F4, F8, FT7, FT8, FC3, FC4, FCZ, T3, T4, C3, C4, CZ, TP7, TP8, CP3, CP4, CPZ, P7, P8, P3, P4, PZ, 01, 02, and OZ) (QuikCap, Neuroscan Inc.) While running participants, the research assistant had an active view of the participant through a camera-monitor set-up, as well as the electrical data being collected through each electrode. Electrode impedance was reduced to less than 5 KOhms before data acquisition. The data was sampled at a rate of 1,000Hz, which was later digitally reduced to 250Hz for analysis. Electrodes were also placed on the left and right temples of the participant (HEOL, HEOR) to record eye movements to the left and the right, and placed on the top and the bottom of the participant’s left eye (VEOU, VEOL) to record eye movements up and down. An electrode was placed on the tip of the participant’s nose in order to serve as a reference.

### Scoring

Scoring responses for the pictures followed the same basic protocol as the rating of the behavioral data. To speed up scoring, subject responses that matched the strings: “nothing,” “none”, “unsure”, and “idk” were automatically marked as a miss.

### EEG Data Analysis

The ERP waveforms analyzed for targets were only included from trials in which a rater marked the participant’s responses as correct. All ERP waveforms regardless of participants’ responses were included for false target analysis. All ERPs were time-locked to the target onset and the false target onset separately, and then epoched in a time range of three seconds (1 second prior to onset, two seconds post). All baseline activity from a window of −200ms to 0ms relative to target onset was removed from each trial. The ERP data was resampled to a rate of 250Hz and bandpass-filtered with a 0.3Hz highpass and a 20Hz lowpass. Analysis was performed using a series of custom scripts utilizing EEGLAB functions.

### Eye movement rejection

The HEOR and HEOL electrodes were used to monitor horizontal eye movements during a trial, and VEOU and VEOL electrodes were used to monitor vertical eye movements. Trials were not included in analysis if a horizontal eye movement voltage magnitude greater than 30uV was detected or any eye movement voltage magnitude greater than 100 uV during the RSVP streams in a moving 32ms time window. All artifact rejection and EEG analysis was performed with a combination of custom MATLAB 2012b scripts and EEGLab 13.3.2b functions^42^.

### Mass univariate analysis

A mass univariate analysis was done on the ERP across all electrodes 200ms to 400ms after target and false target onset, which encompasses the estimated time that an N2pc would appear (~250ms). This was done using the Mass Univariate ERP Toolbox v1.24 (link: http://www.openwetware.org/wiki/Mass_Univariate_ERP_Toolbox). This analysis calculates a t-score for every time point at each electrode, excludes those that do not exceed an uncorrected p-value of .05, and then clusters the remaining t-scores by spatial and temporal proximity. Each cluster’s t-scores are then summed up to calculate the cluster mass. These cluster masses are then compared against randomly generated cluster mass permutations in order to determine the statistical significance of each cluster and control for family-wise error.

### Window-based analysis

A window-based analysis of the ERP data was performed as a confirmatory form of analysis. This analysis uses a sliding-window permutation test to calculate significant differences between set conditions, e.g., targets compared to false targets. The differences between these conditions are then randomized 10,000 times, creating a null hypothesis distribution for each time window across a time range. For the comparison of targets to false targets, this is done using 20ms time steps from 200ms to 400ms. These null distributions are then compared against the true subtracted difference in order to generate a p-value for each time step. The time window is considered significant if it contains at least four steps in a row of a p-value less than 0.05.

## Data availability

All scripts, stimuli, and datasets are provided publicly via repositories on the Open Science Framework (OSF) website, linked below.

Experiment 1:

Preregistration: https://osf.io/gdzxv/

Data and Scripts: https://osf.io/qbnxt/7viewOnly=lf76b4f2507d406d8b4f047a65affdc2

Experiment 2:

Preregistration: https://osf.io/84rvy/

Data and Scripts: https://osf.io/aqyh7/7viewOnly=045d9691138f41c6b5cda07cbf74aade

## Supplemental

To determine if the larger set size in the best-8 categories condition was a factor in the significant difference seen in the N2pc amplitudes, a bootstrapped analysis was performed which resampled the best-8 data to an equal set size as the worst-8’s. The best-8 data was resampled 10,000 times, with the mean amplitude for the N2pc time window calculated each time. These mean amplitude values were then compared against the mean amplitude of the worst-8 dataset (x□=-0.36). Only 2.9% of these values in this nonparametric test equaled or were less than −0.36 value, suggesting that the difference between the best-8 and the worst-8 categories is not an artifact of differences in trial counts. (Supplemental Figure 1)

**Supplemental Figure 1.**
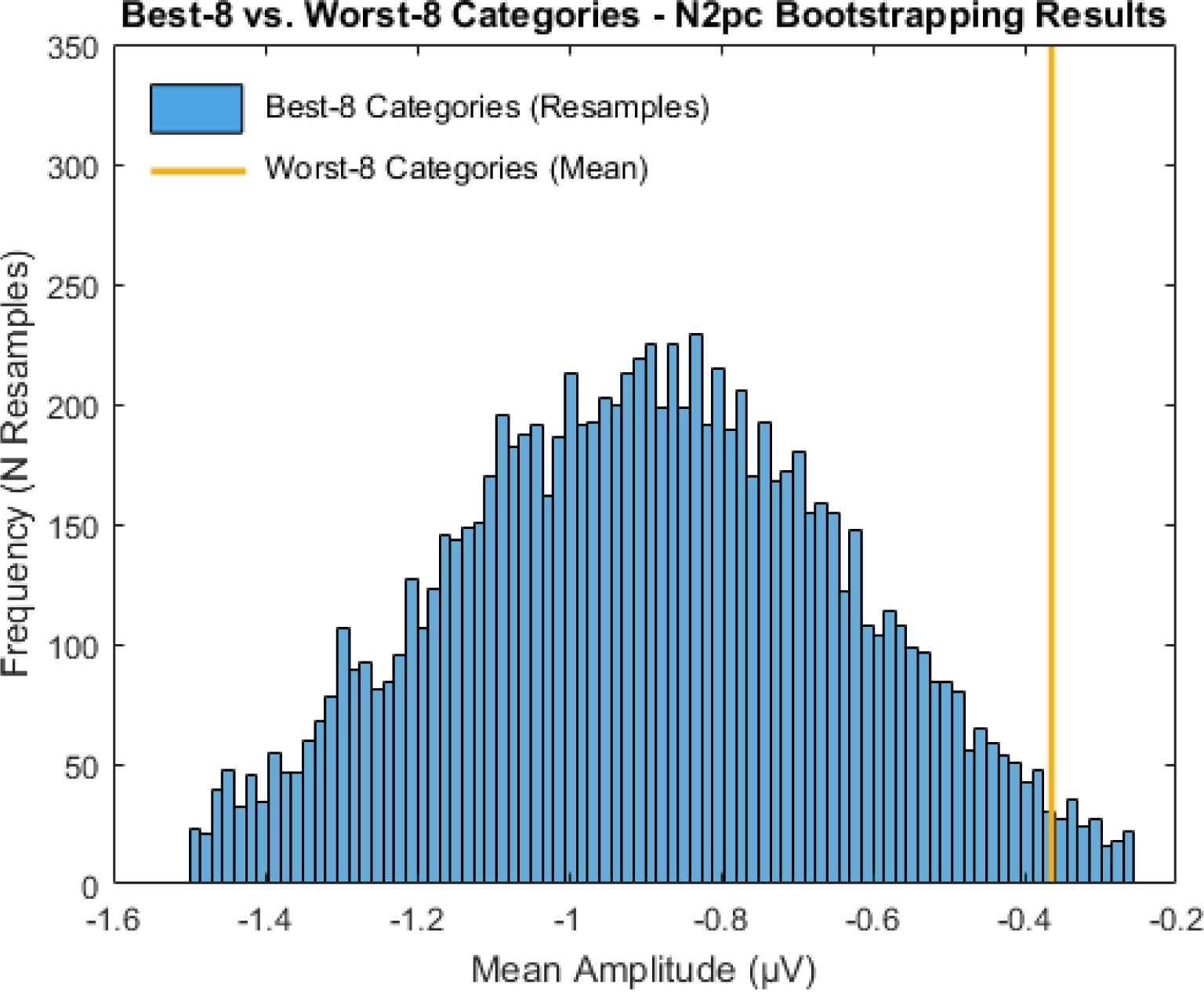
Bootstrapping results comparing resamples of the best-8 categories condition mean amplitude data against the mean amplitude of the worst-8 categories. Only 289 of the 10,000 resamples of the best-8 data had mean amplitudes with an equal or lesser amplitude value than the worst-8 mean (x□=-0.36).

